# Dichloroacetate improves systemic energy balance and feeding behavior during sepsis

**DOI:** 10.1101/2021.07.21.453238

**Authors:** Tae Seok Oh, Manal Zabalwi, Shalini Jain, David Long, Peter W Stacpoole, Charles E McCall, Matthew A Quinn

## Abstract

Sepsis is a life-threatening organ dysfunction by dysregulated host response to an infection. The metabolic aberrations associated with sepsis underly an acute and organism wide hyper-inflammatory response and multiple organ dysfunction; however, crosstalk between systemic metabolomic alterations and metabolic reprograming at organ levels remains unknown. We analyzed substrate utilization by the respiratory exchange ratio, energy expenditure, metabolomic screening and transcriptional profiling in a cecal ligation and puncture (CLP) model, to show that sepsis increases circulating free fatty acids and acylcarnitines but decreases levels of amino acids and carbohydrates leading to a drastic shift in systemic fuel preference. Comparative analysis of previously published metabolomics from septic liver indicates a positive correlation with hepatic and plasma metabolites during sepsis. In particular, glycine deficiency was a common abnormality of both plasma and the liver during sepsis. Interrogation of the hepatic transcriptome in septic mice suggests that the septic liver may contribute to systemic glycine deficiency by downregulating genes involved in glycine synthesis. Interestingly, intraperitoneal injection of the pyruvate dehydrogenase kinase (PDK) inhibitor dichloroacetate (DCA) reverses sepsis-induced anorexia, energy imbalance, dyslipidemia, hypoglycemia, and glycine deficiency. Collectively, our data indicate that PDK inhibition rescues systemic energy imbalance and metabolic dysfunction in sepsis partly through restoration of hepatic fuel metabolism.

## Introduction

Sepsis is a life-threatening condition associated with dysregulated host inflammatory and immune responses to an infection, resulting in immunometabolic suppression, organ failure and high mortality (1–11). However, there is limited knowledge concerning the metabolic alterations in non-immune cells from vital organs in response to sepsis. Metabolic tissues, such as the liver and muscle metabolically reprogram during sepsis with significant consequences for host survival (12–14), indicating systemic metabolism may be subject to dysregulation during severe infection. Therefore, it is important to understand the connection between metabolic and bioenergetic processes and their contribution to the pathogenesis and/or resolution of sepsis.

Metabolomic research in sepsis has provided an atlas of mortality-associated metabolites (15–18). However, knowledge regarding crosstalk between systemic metabolic profiles and intra-organ metabolic reprogramming remains scattered and without clarification of cause and effect in response to sepsis. Indeed, circulating carbohydrates and other small organic molecules and fatty acids are significantly altered in sepsis (19–23), although the underlying mechanisms accounting for these changes and their clinical significance are uncertain, as are their potential as therapeutic targets. Accordingly, we undertook a comprehensive metabolomic screening and energy expenditure of the consequences of sepsis in a murine model of cecal ligation and puncture (CLP) to gain insight into the systemic metabolic manifestations of chronic sepsis. Furthermore, we undertook a comparative analysis of previously published hepatic metabolomics during sepsis to shed light on the potential role of the liver in promoting the systemic metabolic phenotype of sepsis (13). Our study focused on the disease tolerance sepsis state as reported (13, 24), and targeted the mitochondrial pyruvate gate for pyruvate oxidation as a potential contributor to systemic metabolic and bioenergetic reprogramming during sepsis,

## Results

### Sepsis shifts fuel utilization and reduces systemic energy expenditure

To investigate the systemic consequences of sepsis, we first measured food intake up to 30 h following induction of cecal ligation puncture (CLP), using metabolic chambers. In line with previous findings (25, 26), food intake was significantly reduced in response to polymicrobial infection compared to the sham control (Fig. 1A). We next performed unbiased global metabolomics on plasma of control and septic mice at 30 h post-CLP in alignment with our previous studies of the tissue tolerance phenotype of sepsis (13, 24) by Ultrahigh Performance Liquid Chromatography-Tandem Mass Spectroscopy (UPLC-MS/MS). Plasma levels of glucose, fructose, pyruvate and lactate were significantly lower in septic vs. control mice, while circulating succinate levels showed a trend toward lower concentrations (p=0.066) (Figs 1B-F)

**Fig. 1.**
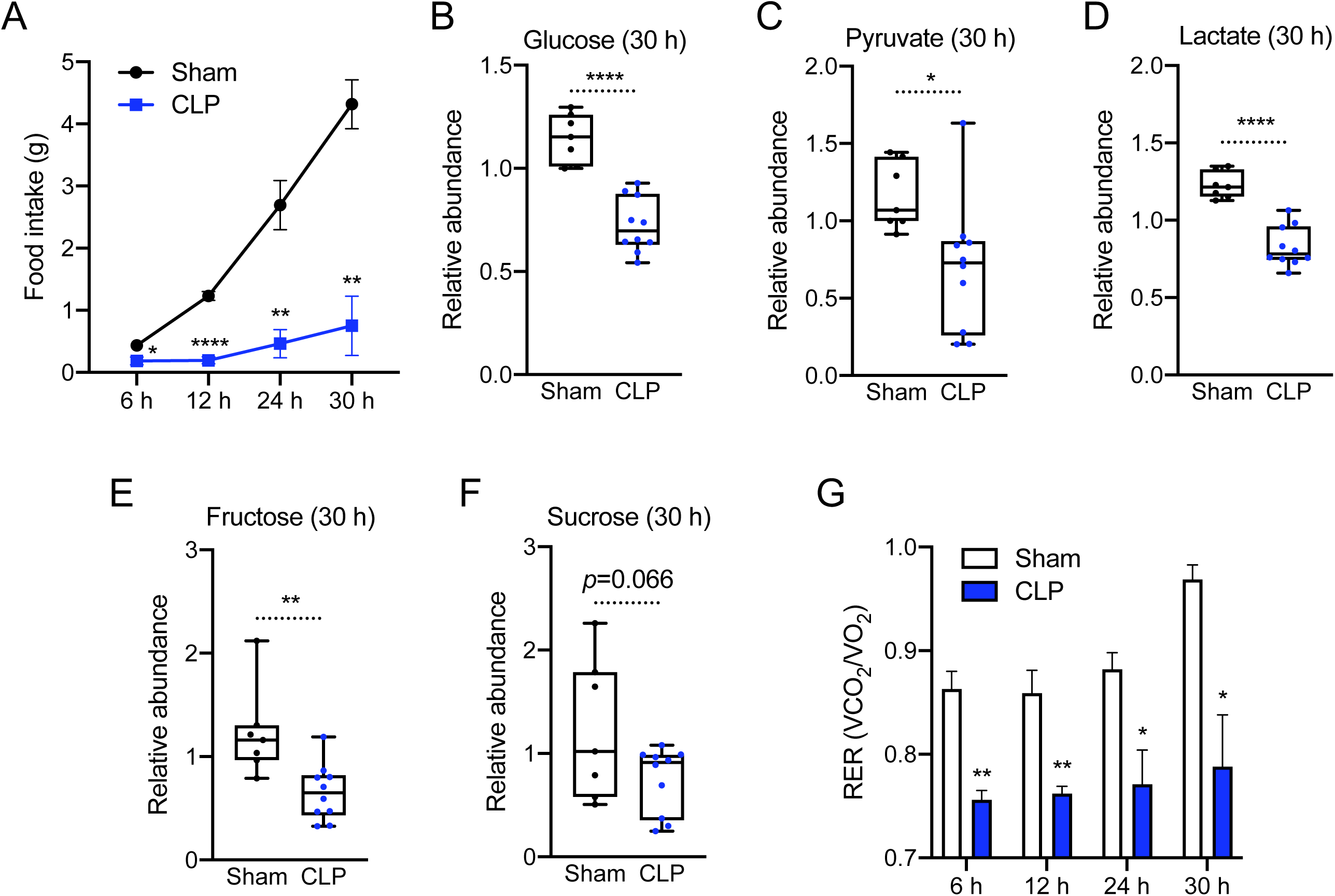
Sepsis reprograms systemic metabolism in response to carbohydrate reduction. *(A)* Cumulative food intake of sham and cecal ligation and puncture (CLP) mice for 30 h (n = 3 sham; 4 CLP). *(B-F)* Relative carbohydrate levels measured by Ultrahigh Performance Liquid Chromatography-Tandem Mass Spectroscopy (UPLC- MS/MS) from plasma of sham and CLP mice 30 h post-surgery (n = 7 sham; 10 CLP). *(G)* Respiratory exchange ratio (RER) of sham and CLP mice at various time points (n = 3 sham; 4 CLP). \**p <* 0.05, \*\**p* < 0.01, \*\*\*\**p* < 0.0001.

We next interrogated the respiratory exchange ratio (RER) to gain further insight into the effects of sepsis on fuel utilization (27, 28). Consistent with the parallel reduction in circulating carbohydrate concentrations, septic mice showed reduced RER values close to 0.7 (Fig. 1G), suggesting a shift from carbohydrates as the primary fuel source to fatty acids, in contrast to controls, in which the RER value was closer to 0.9. (Fig. 1G). We next compared energy expenditure adjusted by body mass of sham and CLP mice. Septic mice showed a significant decrease in energy expenditure regardless of body weight (Fig. 2A) in addition to a trend (*p*=0.057) of less oxygen consumption (Fig. 2B). These data suggest that sepsis induces a state of anorexia that is associated with a shift from carbohydrate to fat as a primary fuel source culminating in impaired systemic metabolism.

**Fig. 2.**
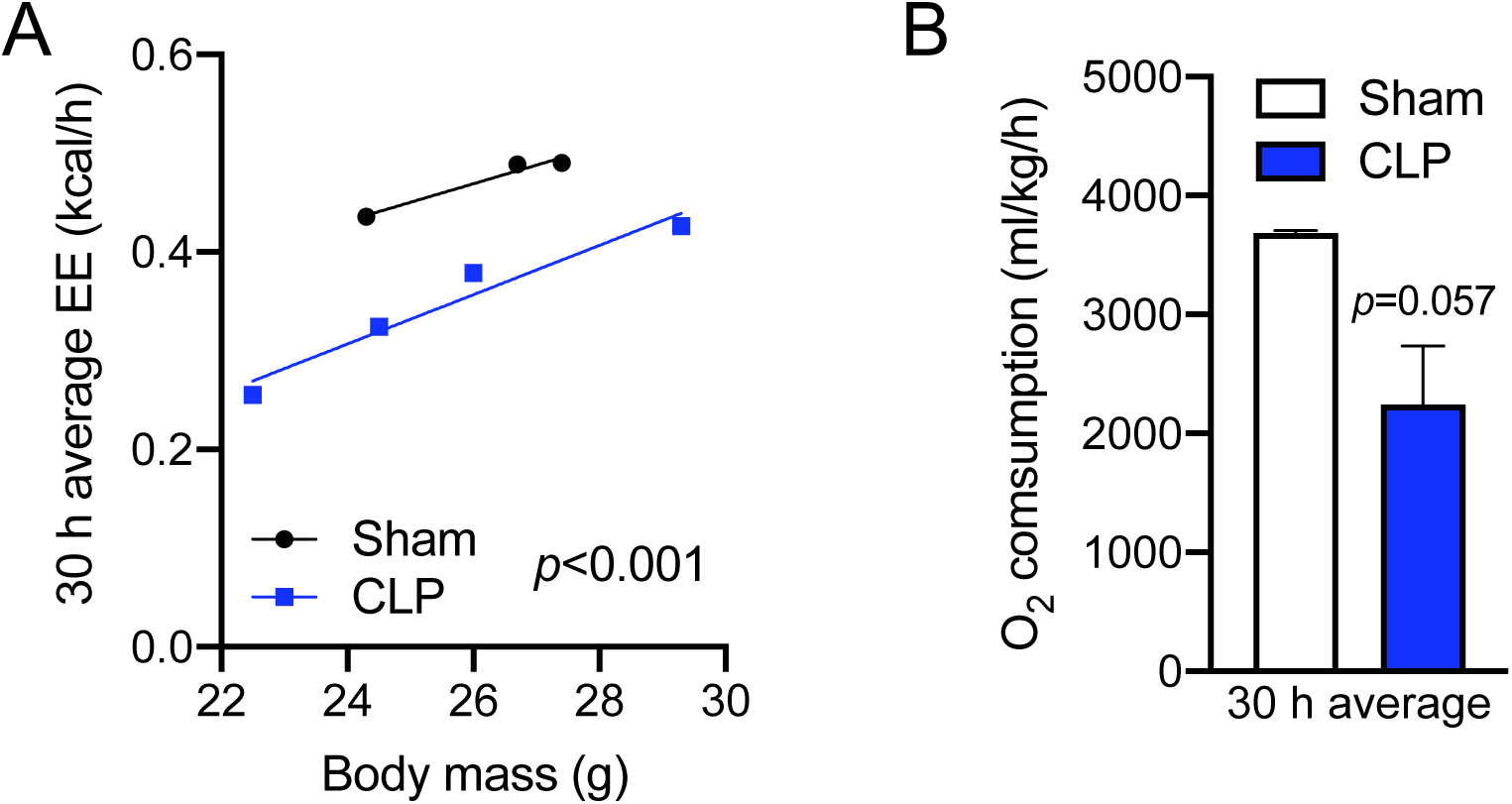
Sepsis impairs systemic energy expenditure. *(A)* Energy expenditure (EE) ANCOVA analysis assessed with body mass of sham and CLP mice (n = 3 sham; 4 CLP). *(B)* O2 consumption measured by metabolic cages for 30 h (n = 3 sham; 4 CLP).

### Sepsis dysregulates circulating lipids and amino acids

In order to further investigate sepsis-induced changes in circulating metabolites contributing to energy imbalance, we evaluated our global unbiased metabolomic screening in plasma from sham and CLP mice. Among 782 metabolites detected, 256 metabolites significantly accumulated (red) and 124 metabolites significantly decreased (green) in septic mice (Fig. 3A). We next performed enrichment analysis using small molecule pathway database (SMPDB) by MetaboAnalyst 5.0 (https://www.metaboanalyst.ca) of those metabolites significantly altered by sepsis. Pathway analysis revealed that perturbations in lipid metabolism, including accumulation of free fatty acids (FFAs), contributed importantly to the plasma metabolomic signature of septic mice, whereas pathways of amino acid metabolism were generally depressed (Fig. 3B-E). Together, these data suggest that sepsis leads to dyslipidemia and depletion of amino acids in the circulation.

**Fig. 3.**
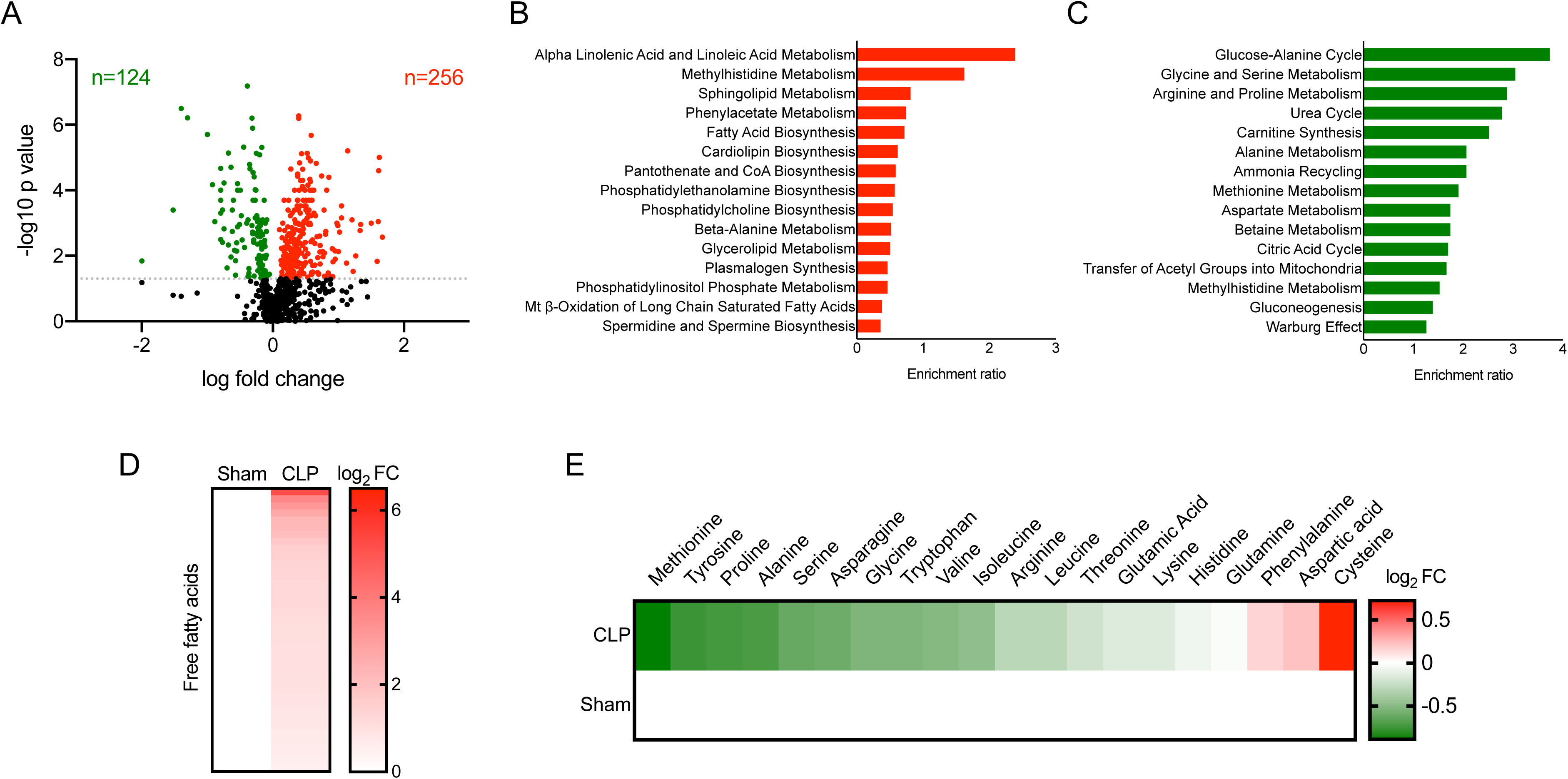
Sepsis elicits dyslipidemia and depletes circulating amino acids and carbohydrates. *(A)* Volcano plot of significantly altered plasma metabolites from sham and CLP mice measured by UPLC-MS/MS (n = 7 sham; 10 CLP). Top 15 metabolic pathways subject to *(B)* accumulated metabolites and *(C)* reduced metabolites in plasma identified by enrichment analysis of sham versus CLP mice (n = 7 sham; 10 CLP). Heatmap depiction of average log2 fold change in plasma *(D)* free fatty acids and *(E)* amino acids in sham and CLP mice 30 h post-surgery measured by UPLC-MS/MS (n = 7 sham; 10 CLP).

### Hepatocytes contribute to systemic dyslipidemia during sepsis

We and others previously reported that sepsis promotes hepatic steatosis in both rodents and humans (13, 29, 30). Given that hepatocytes are involved in the dynamics of circulating free fatty acids (FFAs) (31), we sought to determine whether there is a connection between hepatic steatosis and systemic dyslipidemia in sepsis. Comparing previously published hepatic metabolomics (13) with our plasma metabolomic screen revealed that 399 plasma and 244 liver metabolites were significantly altered in experimental sepsis. Among those metabolites, 72 showed overlap between plasma and hepatocytes (Fig. 4A) and this relationship was positively correlated (Fig. 4B). In addition, those plasma and hepatic metabolites that accumulated in sepsis (Fig. 4B) were enriched in fatty acids (Fig. 4C) as previously reported in humans (22).

**Fig. 4.**
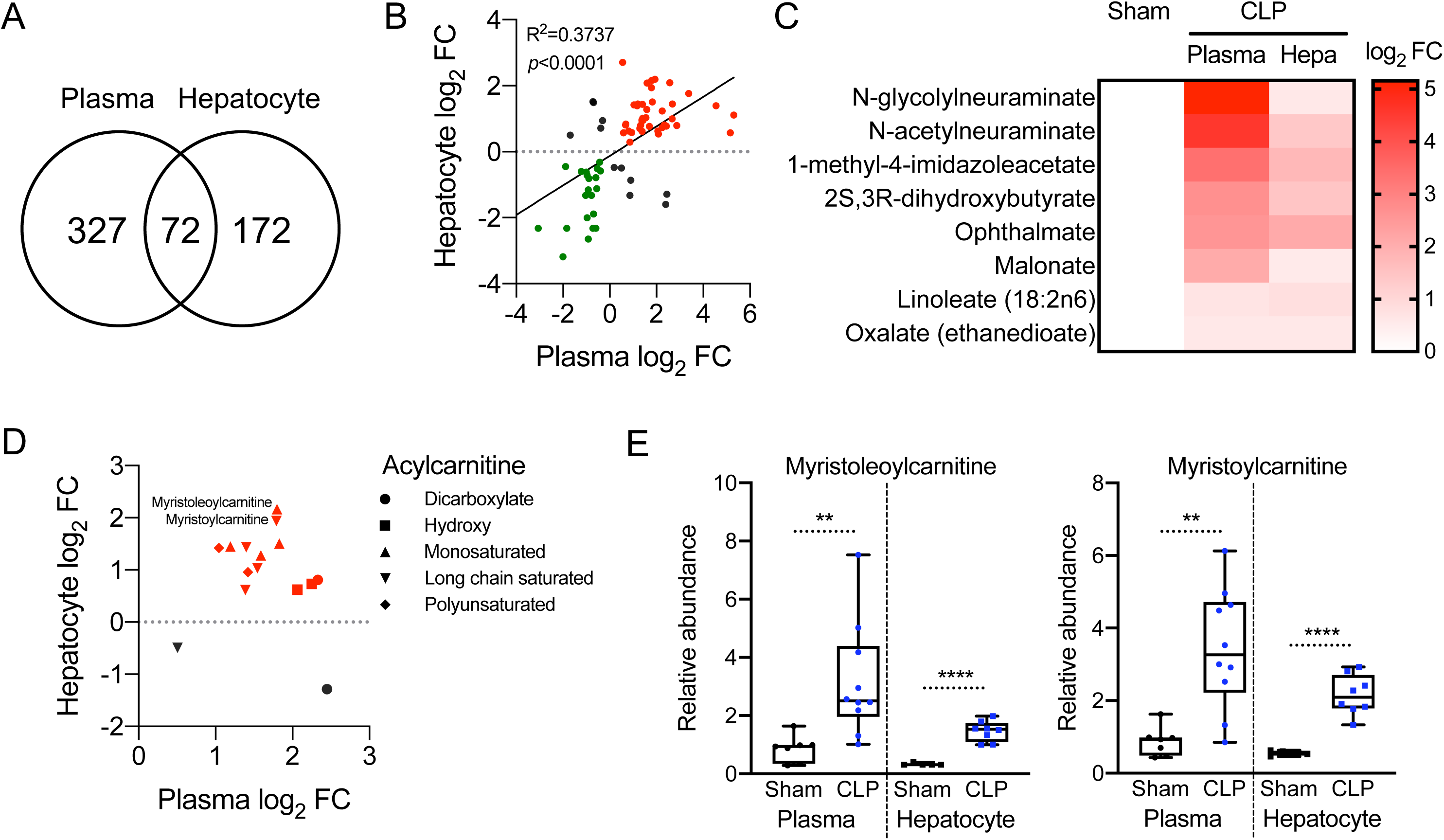
Systemic dyslipidemia correlates with hepatic steatosis during sepsis. *(A)* Venn diagram of significantly altered metabolites from plasma and isolated hepatocytes in sham and CLP mice 30 h post-surgery measured by UPLC-MS/MS (n = 7 sham; 10 CLP for plasma, n = 5 sham; 8 CLP for hepatocytes). *(B)* Correlation analysis with overlapping metabolites between plasma and isolated hepatocytes in sham and CLP mice 30 h post-surgery measured by UPLC-MS/MS (n = 7 sham; 10 CLP for plasma, n = 5 sham; 8 CLP for hepatocytes). Red and green indicate fold increases and decreases in both plasma and hepatocytes, respectively. *(C)* Heatmap depiction of average log2 fold change in fatty acids showing positive correlation between plasma and isolated hepatocytes in sham and CLP mice 30 h post-surgery measured by UPLC-MS/MS (n = 7 sham; 10 CLP). *(D)* Distribution of various acylcarnitine species plotted by average fold change in plasma and isolated hepatocytes in sham and CLP mice 30 h post-surgery measured by UPLC-MS/MS (n = 7 sham; 10 CLP for plasma, n = 5 sham; 8 CLP for hepatocytes). *(E)* Representative acylcarnitine species significantly accumulated in plasma and isolated hepatocytes in CLP mice compared to sham mice 30 h post-surgery measured by UPLC-MS/MS (n = 7 sham; 10 CLP for plasma, n = 5 sham; 8 CLP for hepatocytes).

Acylcarnitines are reported to rewire stress-response inflammatory signaling mediators in monocytes, inducing the secretion of inflammatory cytokines and chemokines (32). Furthermore, starvation leads to increased concentrations of serum and liver acylcarnitines (33). Increases in acylcarnitine species represented another class of metabolites that accumulated in both plasma and liver (Fig. 4D). Two representative acylcarnitines, myristoleoylcarnitine and myristoylcarnitine, showed significant increases in both plasma and hepatocytes (Fig. 4*E*), consistent with the notion that hepatic and/or adipose tissue acylcarnitines may contribute to the systemic dyslipidemia of sepsis.

### Sepsis dysregulates glycine metabolism

We previously reported that mouse septic hepatocytes have low levels of glycine (13), consistent with decreases in plasma glycine levels (Fig. 2E). Glycine is synthesized from serine, threonine, choline and hydroxyproline via inter-organ metabolism, primarily by the liver and kidneys (34). In order to investigate a possible hepatic contribution to systemic glycine deficiency in sepsis, we re-evaluated our previously published RNA-seq gene expression data with regard to enzymes involved in glycine metabolism in livers 30 h post-sepsis (GSE167127) (13). Eight genes involved in the regulation of glycine levels significantly decreased, including glycine N-methyltransferase (*Gnmt*), serine hydroxymethyltransferase 1 (*Shmt1*), *Shmt2*, caspase 7 (*Casp7*), sarcosine dehydrogenase (*Sardh*), glycine C-acetyltransferase (*Gcat*), 5’-aminolevulinate synthase 1 (*Alas1*), and glycine-N-acyltransferase (*Glyat*) (Fig. 5A).

**Fig. 5.**
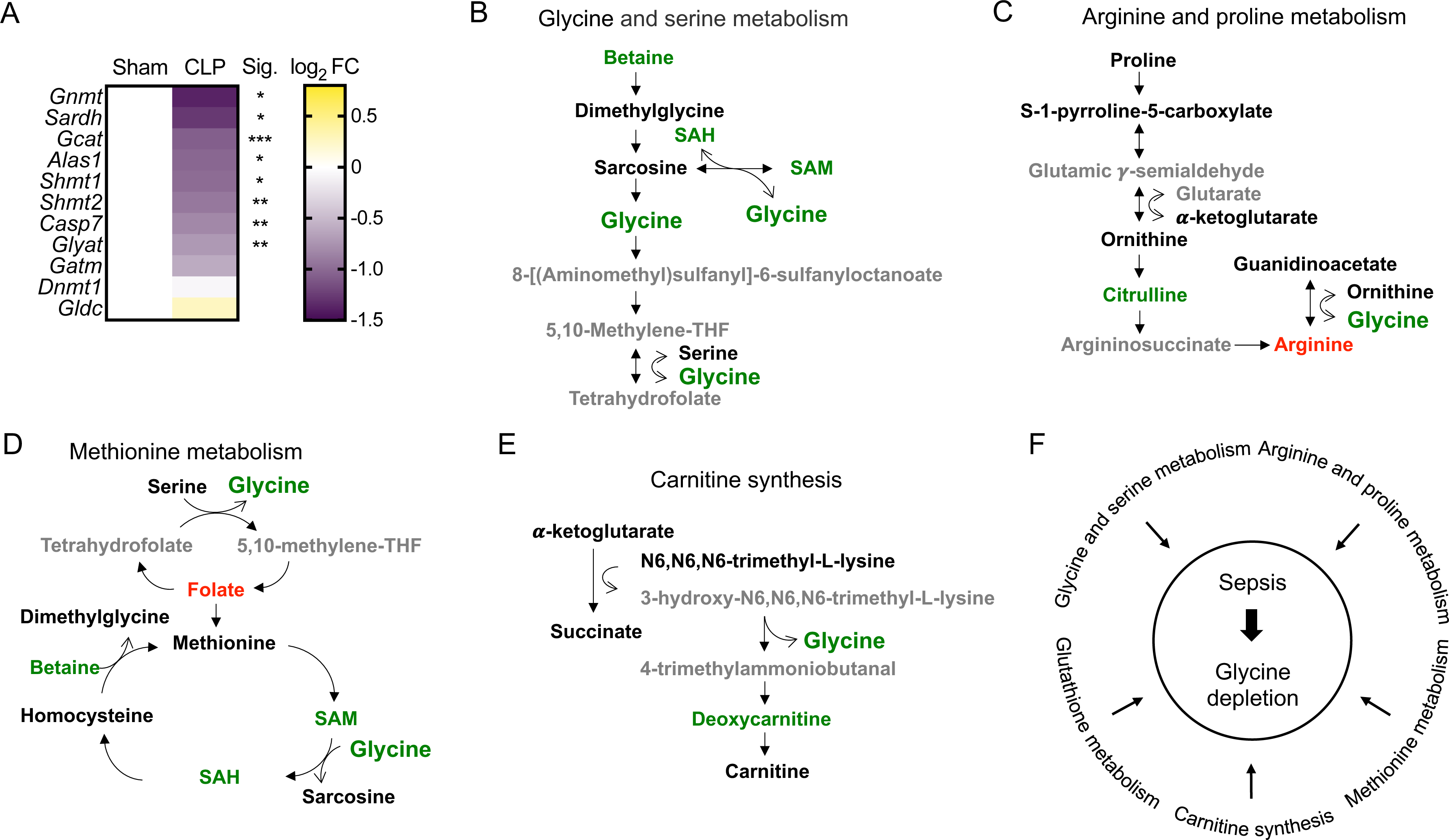
Sepsis dysregulates glycine metabolism. *(A)* Heatmap depiction of average log2 fold change in gene expression involved in glycine metabolism assessed by RNA-seq in sham, CLP, and CLP+DCA 30 h post-surgery (n = 4 mice per group). *(B-E)* Schematic representation of hepatic metabolites contributing to glycine depletion during chronic sepsis. Red denotes a metabolite increased in response to sepsis; green indicates a metabolite decreased in response to sepsis; black indicates a metabolite unchanged in response to sepsis; grey indicates a metabolite not measured in our metabolomic screening. *(F)* Metabolic pathways leading to glycine depletion during sepsis.

In line with altered transcriptional regulation of these genes, we found that sepsis changed the relative abundance of metabolites that lead to glycine deficiency. In particular, betaine, S-adenosyl-L-homocysteine (SAH) and S-adenosylmethionine (SAM) were reduced in the glycine and serine metabolic pathway (Fig. 5B). GNMT reversibly converts sarcosine and SAH to glycine and SAM (35). Downregulation of *Gnmt* gene expression during sepsis was the most robust among those genes involved in glycine metabolism (Fig. 5A). Sepsis also reduced hepatic citrulline concomitant with a shift in hepatic arginine in the arginine and proline metabolic pathway (Fig. 5C), suggesting glycine may be utilized as substrates to support arginine synthesis as an alternative/additional mechanism contributing to glycine deficiency.

Folate accumulated in septic livers as a metabolite in the methionine metabolic pathway (Fig. 5D). SHMT converts tetrahydrofolate and serine to 5,10-methylene-THF and glycine (36) and this component of one carbon metabolism was significantly downregulated in septic livers (Fig. 5A), possibly contributing to glycine deficiency. In contrast, metabolites in the carnitine synthetic pathway did not show distinct alterations, and SHMT also produces glycine as a side product using 3-hydroxy-N6, N6,N6-trimethyl-L-lysine and 4-trimethylammoniobutanal as a substrate and a resultant product, respectively (Fig. 5E).

We also previously reported that sepsis depletes hepatic glutathione levels (13), which may be a consequence of glycine deficiency. These alterations in hepatic metabolites also aligned with changes in circulating metabolites (Fig. S1). Taken together, multiple metabolic pathways that affect glycine levels are modulated by sepsis (Fig. 5F).

### DCA restores feeding behavior and fuel utilization

We have previously reported that pyruvate dehydrogenase kinase (PDK) inhibition by dichloroacetate (DCA) rebalances immunometabolic and mitochondrial respiration and anabolic energetics during sepsis, thereby increasing survival in septic mice (24). Furthermore, DCA reverses dysregulated hepatocyte metabolism, including triglyceride accumulation and mitochondrial dysfunction (13). Given that DCA restores the hepatic metabolome and increases hepatic mitochondrial energetics to promote survival in septic mice, we hypothesized that PDK inhibition might also reverse sepsis-induced anorexia. To assess this, we housed septic mice treated with or without DCA in metabolic cages after sepsis onset. Strikingly, DCA administration completely normalized food intake in CLP mice when administered one hour after CLP (Fig. 6A). DCA treatment also rebalanced abnormal concentrations of circulating carbohydrates (Fig. 6B) and free fatty acids (FFAs) (Fig. 6C). Together, these data support that stimulating PDC by reducing function of its principle inactivator, PDK, may restore systemic fuel availability during sepsis.

**Fig. 6.**
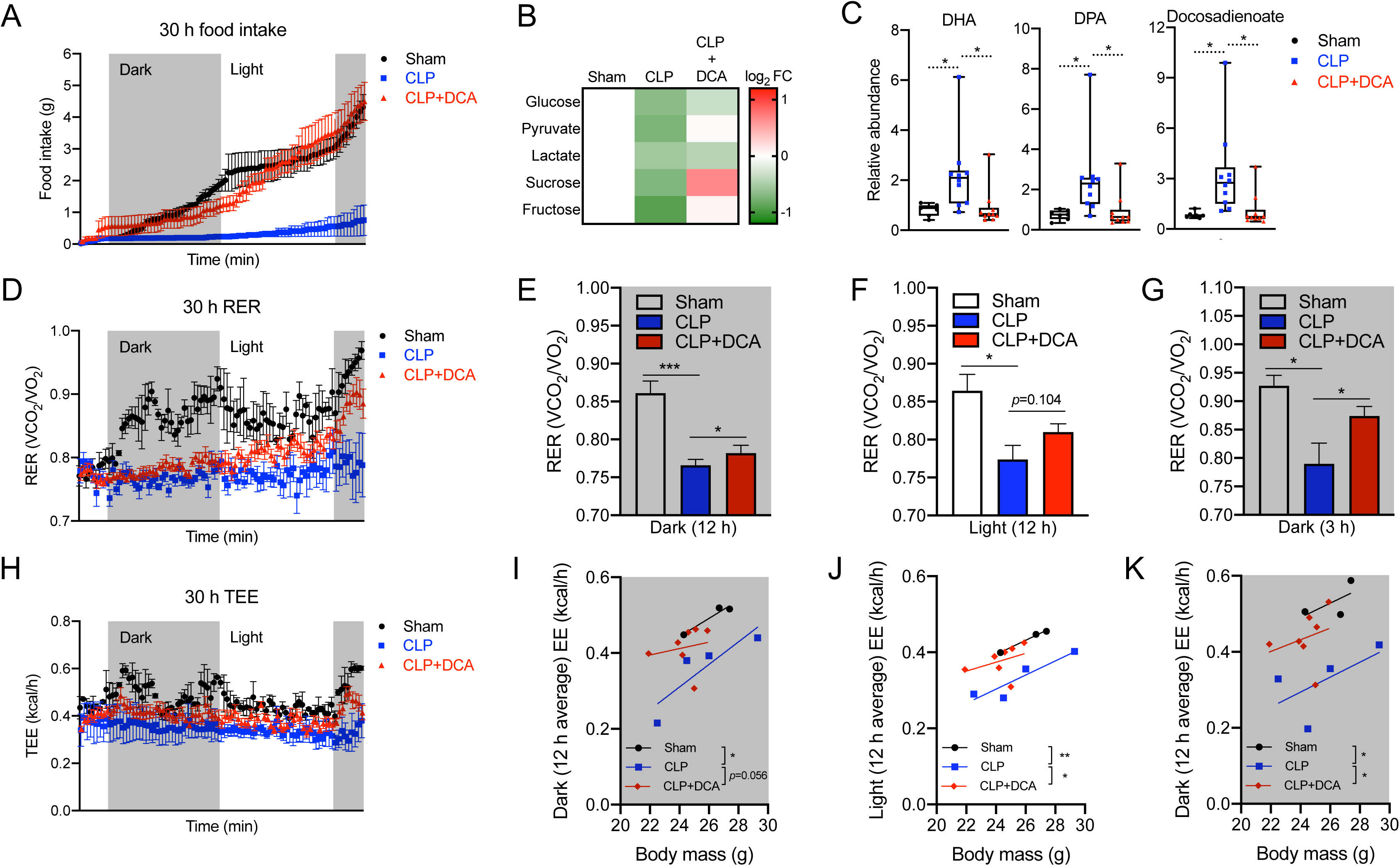
DCA recovers sepsis-induced anorexia and reprograms systemic fuel utilization. *(A)* Cumulative food intake of sham, CLP, and CLP+DCA over 30 h (n = 3 sham; 4 CLP; 7 CLP+DCA). *(B)* Heatmap depiction of average log2 fold change in carbohydrate levels measured by UPLC-MS/MS from plasma of sham, CLP, and CLP+DCA 30 h post-surgery (n = 3 sham; 4 CLP; 7 CLP+DCA). *(C)* Relative fatty acid levels measured by UPLC-MS/MS from plasma of sham, CLP, CLP+DCA 30 h post-surgery (n = 7 sham; 10 CLP; 10 CLP+DCA). *(D)* RER of sham, CLP, and CLP+DCA for 30 h (n = 3 sham; 4 CLP; 7 CLP+DCA). Average RER for *(E)* 12 h dark cycle, *(F)* 12 h light cycle, and *(G)* 3 h dark cycle. (*H*) Total energy expenditure (TEE) of sham, CLP, and CLP+DCA for 30 h (n = 3 sham; 4 CLP; 7 CLP+DCA). Average EE for *(I)* 12 h dark cycle, *(J)* 12 h light cycle, and *(K)* 3 h dark cycle. \**p <* 0.05, \*\**p* < 0.01, \*\*\**p* < 0.001.

To confirm that metabolite rewiring might lead to changes in systemic energetics, we assessed DCA’s effects on RER and EE. Decreased RER in septic mice was reversed by DCA in a time-dependent manner throughout 30 h post CLP (Fig. 6D). Of note, RER in the dark (feeding) cycles showed significant recovery by DCA (Fig. 6E and G) with a modest trend (*p*=0.104) during the light cycles (Fig. 6F), compatible with circadian rhythm potentially affecting sepsis mortality (37). Reduced TEE was also rescued by DCA during sepsis (Fig. 6H). ANCOVA analysis assessed with body mass showed that the 12 h average EE in the first dark cycle elicited a strong trend (*p*=0.056) of restoration by DCA during sepsis (Fig. 6I). The same analysis in the light cycle and the second dark cycle exhibited significant restoration by DCA during sepsis (Fig. 6J and *K*). Collectively, these data suggest that PDK inhibition ameliorates sepsis by decreasing anorexia and reprograming systemic fuel utilization.

### DCA ameliorates acylcarnitine accumulation during sepsis

Given that DCA may reverse the switch from carbohydrate to FFA utilization during sepsis, we next evaluated its effects on acylation of carnitines. This was done in consideration for their role in balancing sugar and lipid metabolism (38) and that liver is a major contributor to systemic acylcarnitine levels (39, 40). Accordingly, we found that hepatic expression of carnitine acyltransferase (*Crat)* in septic mice was significantly upregulated and reversed by DCA (Fig. 7A). In accordance with altered *Crat* expression, carnitine levels in the circulation were significantly reduced during sepsis and this reduction was also restored by DCA (Fig. 7B). However, hepatic carnitine levels remained unaltered during sepsis, possibly due to compensatory biosynthesis (Fig. 7B). The rate limiting function of gamma-butyrobetaine dioxygenase (BBOX1) catalyzes mature carnitine from its precursor deoxycarnitine. *Bbox1* mRNA levels showed a decreasing trend (*p*=0.0559) in septic livers (Fig. 7C), and both circulating and hepatic deoxycarnitine levels significantly decreased in septic mice. DCA restored only hepatic deoxycarnitine levels (Fig. 7D). Considering deoxycarnitine is converted to carnitine by BBOX1, hyperactivation of BBOX1 could occur and contribute to high flux of conversion from deoxycarnintine to carnitine in the septic liver. As *Crat* expression implied, sepsis led to acylcarnitine accumulation in hepatocytes as well as in the circulation and DCA reversed this phenomenon (Fig. 7E). In sum, these data suggest that DCA normalizes excessive acylcarnitine contents in the circulation and the liver, regulating hepatic gene expression involved in acylation of carnitines.

**Fig. 7.**
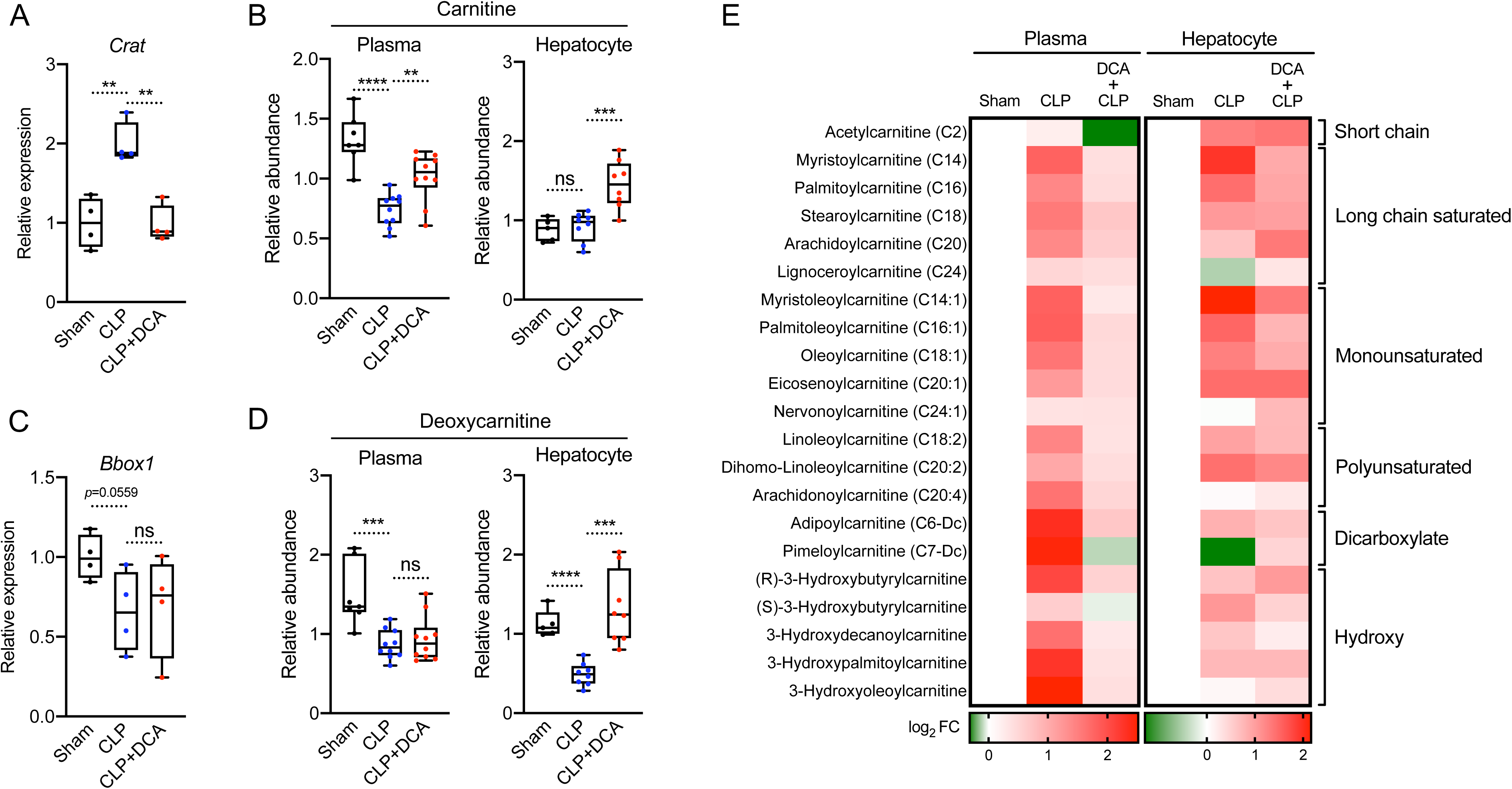
DCA reduces acylcarnitines during sepsis. *(A)* Relative gene expression of carnitine acyltransferase (*Crat*) assessed by RNA-seq in sham, CLP, and CLP+DCA 30 h post-surgery (n = 4 mice per group). *(B)* Relative carnitine levels measured by UPLC- MS/MS from plasma of sham, CLP, CLP+DCA 30 h post-surgery (n = 7 sham; 10 CLP; 10 CLP+DCA). *(C)* Relative gene expression of *Bbox1* assessed by RNA-seq in sham, CLP, and CLP+DCA 30 h post-surgery (n = 4 mice per group). *(D)* Relative deoxycarnitine levels measured by UPLC-MS/MS from plasma of sham, CLP, CLP+DCA 30 h post- surgery (n = 7 sham; 10 CLP; 10 CLP+DCA). *(E)* Heatmap depiction of average log2 fold change in acylcarnitine levels measured by UPLC-MS/MS from plasma of sham, CLP, and CLP+DCA 30 h post-surgery (n = 3 sham; 4 CLP; 7 CLP+DCA). \*\**p* < 0.01, \*\*\**p* < 0.001, \*\*\*\**p* < 0.0001.

### DCA reinstates sepsis-induced glycine deficiency

Given that sepsis dysregulates glycine metabolism in mice, we examined whether DCA might blunt glycine deficiency during sepsis by assessing *GNMT, SARDH, SHMT1* and *SHMT2*, the key enzymes involved in glycine metabolism (Fig. S2). Significant downregulation of these hepatic genes during sepsis reversed after DCA administration (Fig. 8A-D). DCA treatment resulted in a trend (*p*=0.077) for reversed glycine levels by DCA in the septic liver and a significant recovery in circulating glycine levels (Fig. 8E). DCA treatment of septic mice also reversed sepsis-regulated metabolites in the glycine pathway in plasma and liver (Fig. 8F). These included S-methylgultathione (glutathione metabolism), citrulline (arginine and proline metabolism), putrescine (methionine metabolism), and creatine (glycine and serine metabolism), implicating these as liver-driven metabolites that might contribute to systemic glycine deficiency during sepsis.

**Fig. 8.**
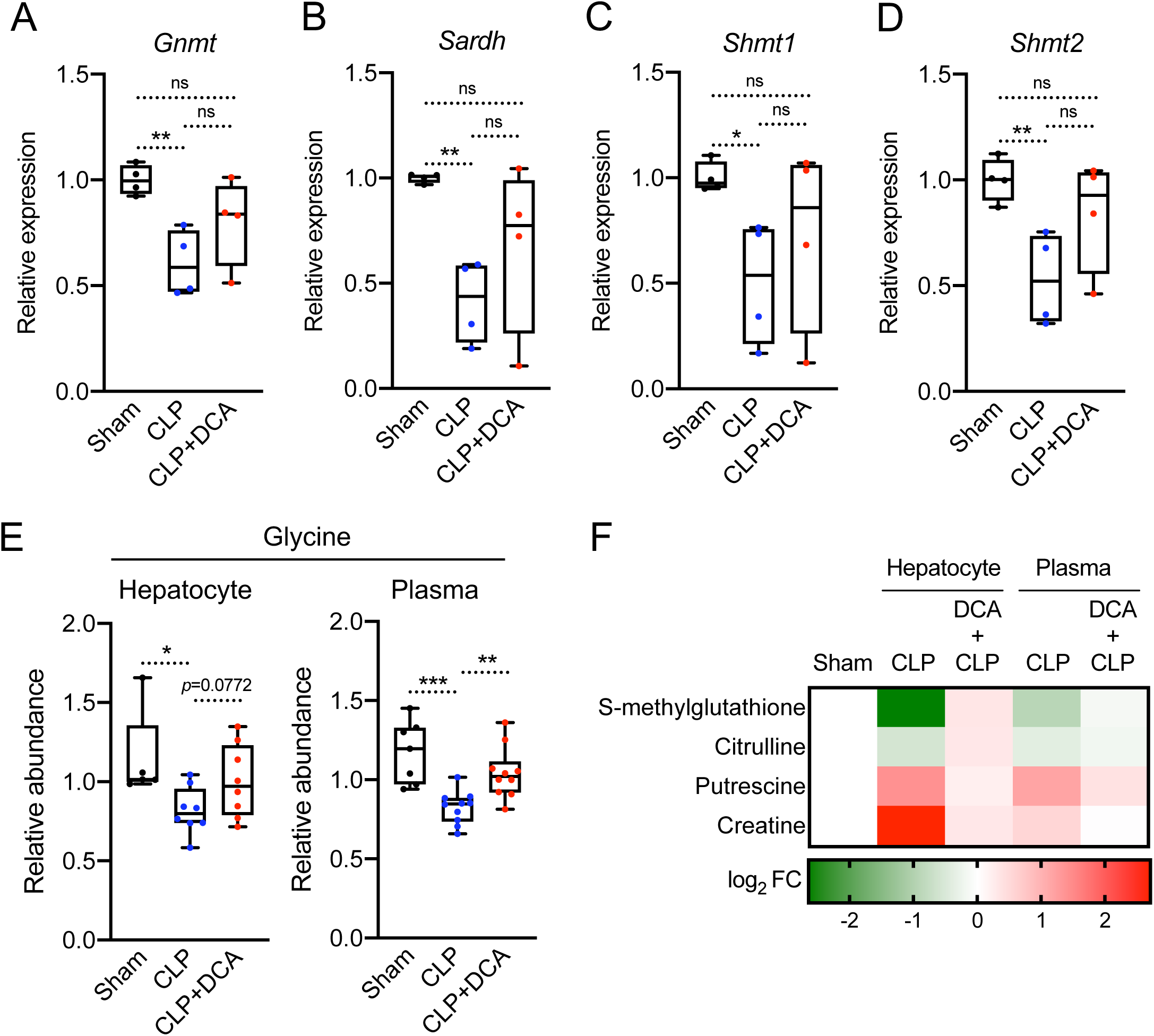
DCA reinstates sepsis-induced metabolic alterations shared by liver and plasma. *(A-D)* Relative gene expression involved in glycine metabolism assessed by RNA-seq in sham, CLP, and CLP+DCA 30 h post-surgery (n = 4 mice per group). *(E)* Relative glycine levels measured by UPLC-MS/MS from plasma of sham, CLP, CLP+DCA 30 h post-surgery (n = 7 sham; 10 CLP; 10 CLP+DCA). *(F)* Heatmap depiction of average log2 fold change in glycine pathway metabolites measured by UPLC-MS/MS from plasma of sham, CLP, and CLP+DCA 30 h post-surgery (n = 3 sham; 4 CLP; 7 CLP+DCA). \**p <* 0.05, \*\**p* < 0.01, \*\*\**p* < 0.001.

## Discussion

Energy metabolism is disrupted during sepsis and is a major contributor to organismal demise (41, 42). Better understanding of interaction and dynamics between systemic metabolic alterations and multiorgan failure provides a logical foundation for therapeutic interventions in sepsis (43). In the present study, we found that sepsis elicits anorexia, alters fuel availability/utilization and, ultimately, culminates in systemic energy imbalance in mice. We identify a possible hepatic contribution of sepsis-induced dyslipdemia and glycine deficiency as a potential target. Our data indicate that administration of the PDK inhibitor DCA reverses dysregulated systemic energy balance and metabolic dysfunction partly through hepatic metabolic regulation, which may ultimately underlie the protective effects of DCA during sepsis (Fig. 9).

**Fig. 9.**
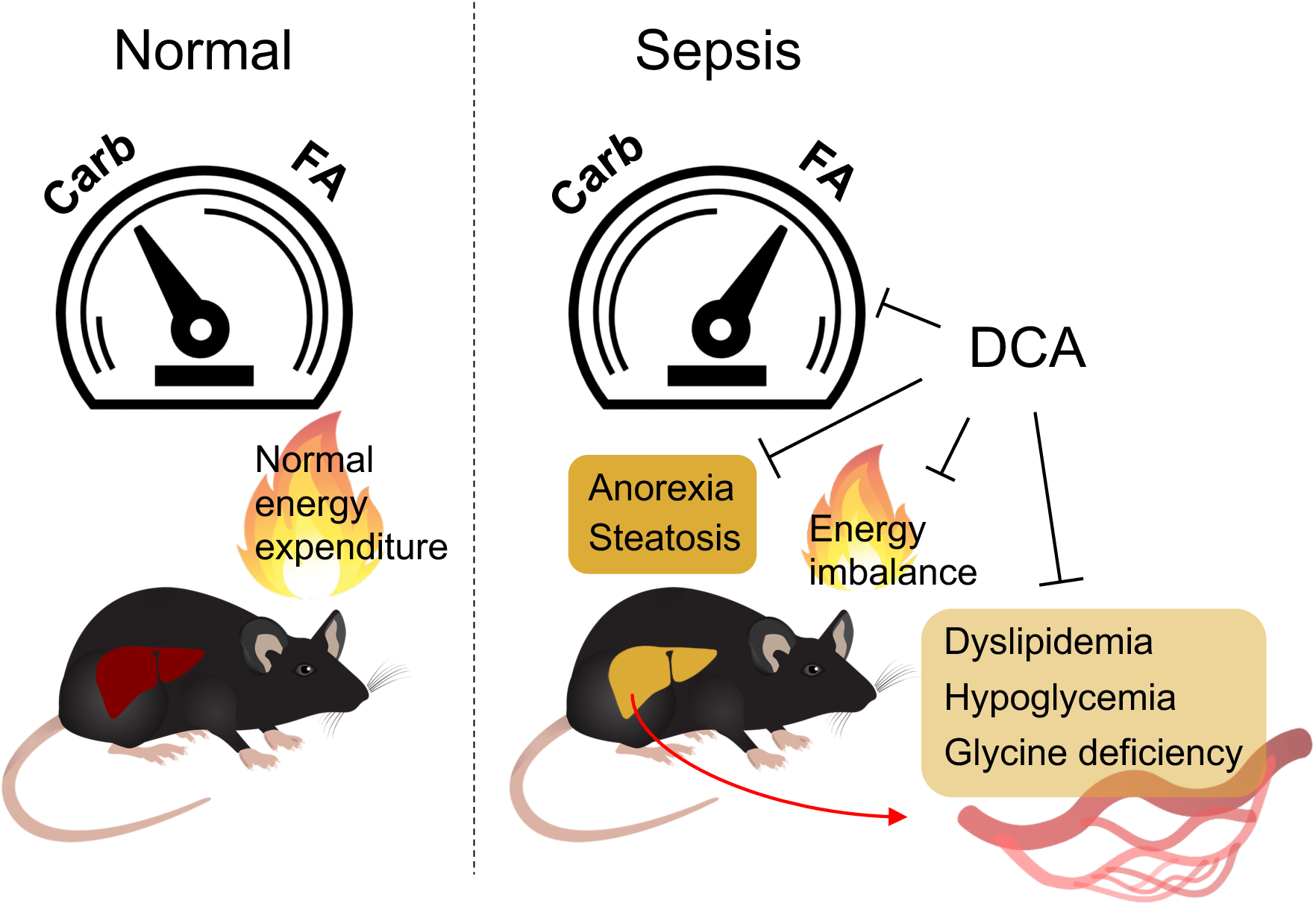
DCA rescues deleterious metabolic alterations during sepsis. Schematic diagram describing detrimental alterations in response to sepsis and reversion effects of DCA.

Fuel utilization during sepsis has been addressed for several decades due to its role in governing alterations of systemic metabolism in critical illness (44–47). Fluctuations in circulating glucose occur commonly in sepsis, and hypoglycemia has been associated inversely with survival (47–49). The murine CLP model also been associated with hypoglycemia in the chronic phase of sepsis (13), which we also observed in the present study. Furthermore, we observed significant increases in circulating free fatty acids during sepsis, which may ultimately underlie the shift in fuel preference from carbohydrate sources to utilization of free fatty acids.

Our metabolomic analysis revealed significant reductions in the circulating levels of both carbohydrates and amino acids in contrast to fatty acids. Others have reported that patients with sepsis have reduced plasma concentrations of most amino acids and higher infusion rates of amino acids are required to maintain plasma concentrations due to increased hepatic extraction of amino acids from plasma (19). In the present study, we found that glycine deficiency is a common abnormality of both plasma and the liver during sepsis, with downregulation of hepatic gene expression for glycine synthesis. Considering glycine as a precursor glutathione and carnitine, glycine deficiency can lead to organ damage from excessive oxidative stress. For example, glycine and cysteine supplementation in HIV-infected patients restored glutathione synthesis and mitochondrial fuel oxidation (48). Although another study suggests that pre-feeding of glycine reduces liver damage and dysregulated systemic inflammatory responses in a rat sepsis model (49), significant gaps exist in our understanding of the putative beneficial effects of glycine during sepsis and additional studies are needed. A potential caveat to interpreting the effect of DCA on glycine levels is that the drug is metabolized to glycine in humans and rodents (50), although the quantitative significance of a single dose of DCA as a contributor to circulating levels of this amino acid under the conditions reported here is likely to have little impact on total intrahepatic or circulating glycine levels.

The carnitine pool, comprised of L-carnitine and its acylated derivatives, is consistently recognized as a prognostic indicator of severe sepsis and septic shock. Increased plasma acylcarnitine predicts high mortality of septic patients (22, 51, 52). In line with these studies, our data suggest a hepatic contribution to systemic acylcarnitine accumulation during sepsis. We show that hepatic *Crat* is highly expressed during sepsis, suggesting that increased acylation of carnitine in the liver contributes to elevated levels of acylcarnitines in plasma. CRAT can drive either a forward acylcarnitine synthesis and its accumulation or reverse it and increase acetyl-CoA regeneration to fuel mitochondrial energetics (53). L-carnitine supplementation in human patients with septic shock elevates serum acylcarnitine levels (54), supporting that acylcarnitine synthesis is active during sepsis.

The mitochondrial pyruvate dehydrogenase megacomplex (PDC) catalyzes the rate-determining step in the aerobic oxidation of glucose and of the glycolytic product pyruvate, to acetyl CoA. Rapid regulation of the complex is mediated mainly by reversible phosphorylation by any one of four pyruvate dehydrogenase kinase (PDK) isoforms and two pyruvate dehydrogenase phosphatase isoforms (55). Pathological up-regulation of PDKs, resulting in inhibition of PDC, has been reported in diverse acquired diseases of metabolic integration and immune dysfunction. We reported that the pan-PDK inhibitor DCA reduces sepsis mortality (24), regulates anabolic and catabolic energy supply in immune cells and hepatocytes (13, 56), as well as reducing TCA cycle tolerance mediator itaconate. Relevant to this is that DCA reverses hepatic metabolic dysfunction and the mitochondrial low energy state in septic mice (13). In the present study, we show that PDK inhibition reverses sepsis-induced anorexia, restores carbohydrate fuel metabolism and rebalances the carnitine:acylcarnitine ratio. Nevertheless, the majority of carnitines are stored in the heart and skeletal muscle (57), and their contribution to carnitine dynamics in liver needs to be considered in the future studies of the metabolic consequences of sepsis. Furthermore, in humans, BBOX1, the essential enzyme for carnitine synthesis, is located in the kidneys and the brain as well as in the liver (58). Therefore, investigating carnitine metabolism among these tissues in response to sepsis would fill an important knowledge gap and might inform one way that DCA improves sepsis survival in mice.

In conclusion, we show that CLP-induced murine sepsis causes major disruption of carbohydrate, fat and amino acid metabolism in both liver and plasma, resulting in significant perturbation of systemic bioenergetics. Pharmacological targeting of the PDC/PDK axis in liver has the potential to overcome sepsis-induced hepatic immunometabolic dysfunction that leads to systemic energy crisis and thus offers a new approach to the development of precision therapeutics for severe sepsis.

## Materials and methods

### Animal Experiments

Animal experiments were conducted as previous described (13). Dichloroacetate (DCA) (Sigma; MO, USA) was administered (25mg/kg) intraperitoneally at various time points post-surgery: for metabolomic screening and RNA-seq DCA was administered 24 hours post-surgery and tissues and plasma were collected 6 hours post DCA administration (30 h post-surgery), for metabolic cages DCA was administered immediately post-surgery.

### Indirect Calorimetry

To measure whole body energy expenditure in live animals, mice were housed individually in metabolic chambers of PhenoMaster (Indirect calorimetry system; TSE systems, Germany) and acclimatized for 3 days with free access to food and water. The Energy expenditure (Oxygen consumption (VO2), Respiratory exchange ratio (RER) and food intake were obtained continuously during a 12 h light and 12 h dark cycle for four days and measured for last 24 h after adaptation.

### Ultrahigh-performance liquid chromatography–tandem mass spectroscopy

Metabolomic screening was performed by previously a described manner (13) with plasma samples from mice described above. Enrichment analysis was performed with small molecule pathway database (SMPDB) by MetaboAnalyst 5.0 (https://www.metaboanalyst.ca). Significantly altered metabolites by sepsis were recognized by human metabolome database (HMDB) ID and used for over representation analysis. Enrichment ratio was computed based on observed hits divided by expected hits in the given pathways.

### Statistical analysis

Unpaired t-test was performed to determine significant relationships between groups in the experiments. Two-way ANOVA was used when comparing more than two groups. Data are represented as mean ± SEM.

## Acknowledgement

This work was supported by R35 GM126922 (CEM). 1-RO1 GM099871 (P.S.).

## Notes

### Competing Interest Statement

The authors have declared no competing interest.

