## Supplementary material for "Dichloroacetate improves systemic energy balance and feeding behavior during sepsis": Suppl Figs

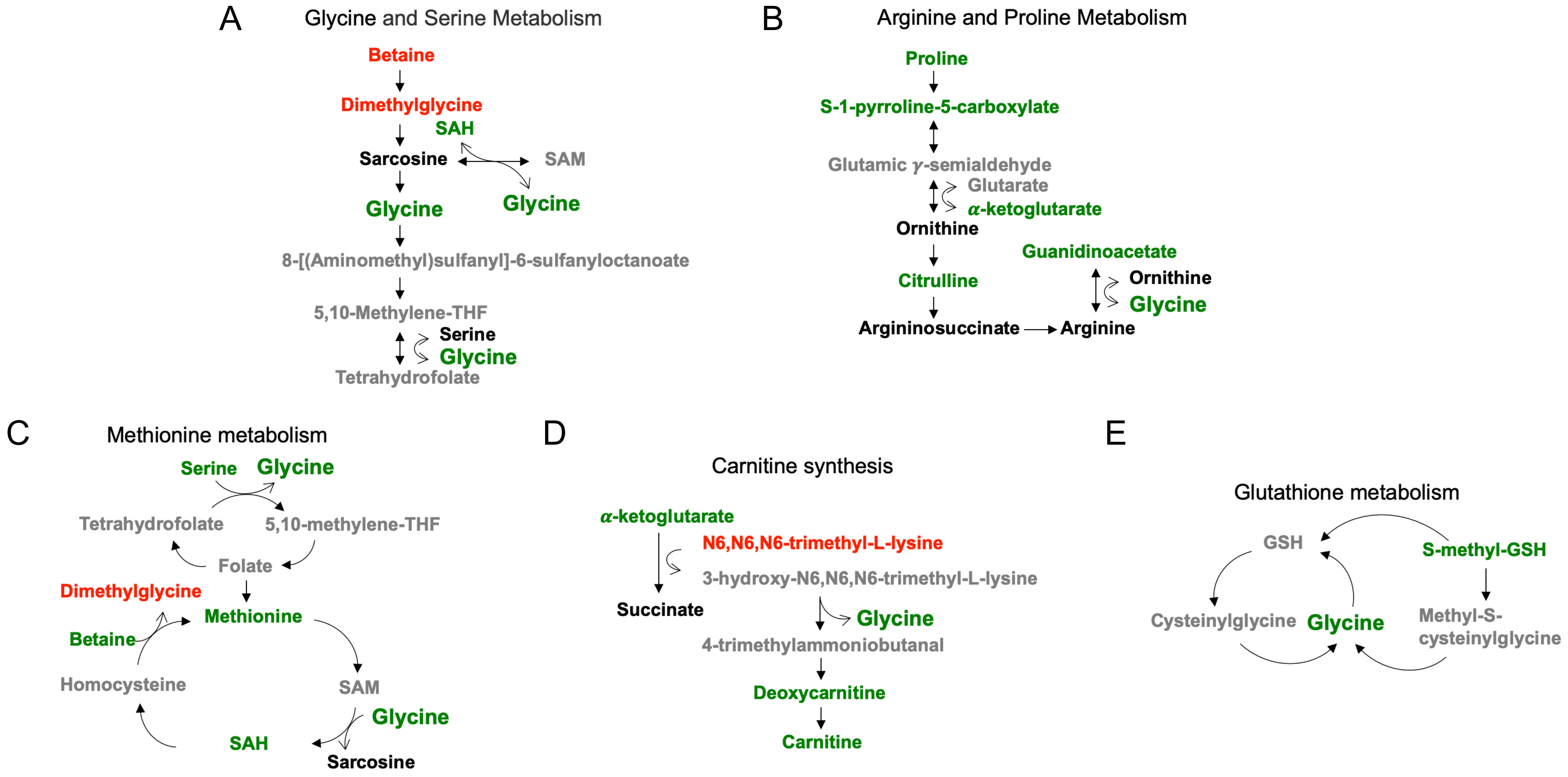


**Fig. S1. Changes in circulating metabolites involved in glycine deficiency in response to sepsis.** *(A-E)* Schematic representation of plasma metabolites contributing to glycine depletion during chronic sepsis. Red denotes a metabolite increased in response to sepsis; green indicates a metabolite decreased in response to sepsis; black indicates a metabolite unchanged in response to sepsis; grey indicates a metabolite not measured in our metabolomic screening.

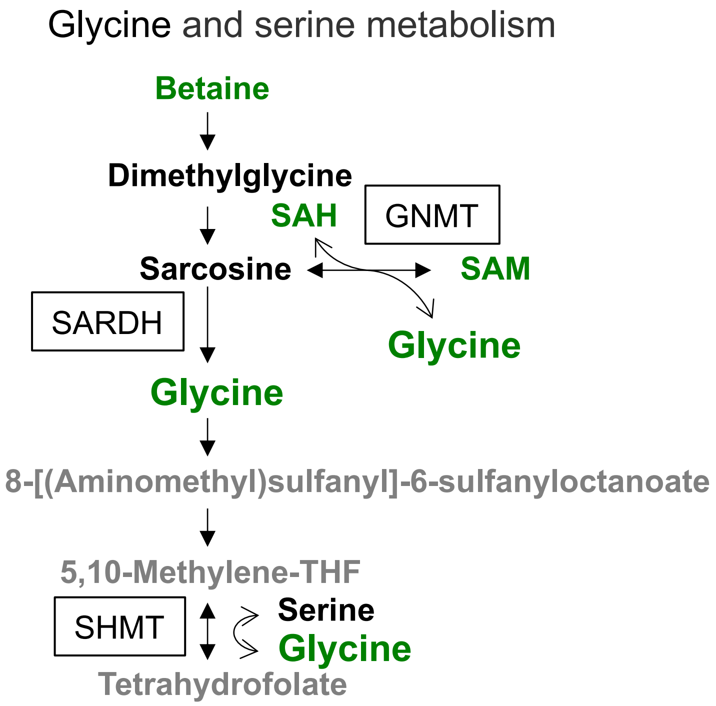


**Fig. S2. Enzymes that lead to glycine deficiency during sepsis.** Enzymes involved in glycine synthesis are shown in rectangles in the glycine and serine metabolic pathway. Green indicates a metabolite decreased in response to sepsis; black indicates a metabolite unchanged in response to sepsis; grey indicates a metabolite not measured in our metabolomic screening.
